# Increased clonality, decreased allele diversity and high genetic structure in tetraploid sea anemone *Aulactinia stella* populations from North Pacific to Atlantic across the Arctic Ocean

**DOI:** 10.1101/2025.04.24.650399

**Authors:** Ekaterina Bocharova, Aleksandr Volkov, Solenn Stoeckel

**Author notes:** Peer-reviewed and recommended by PCI Evolutionary Biology. The full recommendation and reviews are available at https://evolbiol.peercommunityin.org/articles/rec?id=866.

## Abstract

Reproductive mode is a key factor shaping genetic diversity, evolutionary potential, and the processes of dispersal and colonization. Clonality is particularly common in harsh environments and at the margins of species ranges, where it supports persistence, enables rapid growth, and promotes the maintenance of locally adapted genotypes. In the rapidly changing Arctic, increasing ecological connectivity is eroding historical barriers for sessile species. Evaluating genetic diversity in this context, before global change further alters Arctic ocean, is essential for understanding evolutionary dynamics during range expansion and for informing conservation strategies.

*Aulactinia stella* is a circumpolar sea anemone with physiological characteristics in laboratory conditions suggesting a potential for clonal reproduction. In this study, we investigated its reproductive modes in natural populations across the Arctic ocean, from the northern Pacific to the Atlantic, and examined how genetic diversity is structured between adults and juveniles at five sampled sites.

Across all study sites, we observed only females or individuals lacking gonads, with the exception of Kamchatka, where males were also present. Genetic indices and changes in genotype frequencies between adults and juveniles confirmed that this species reproduces partially by parthenogenesis. Populations on the Atlantic side were highly clonal with clonal rates (*c*) estimated at 80-99%, whereas populations on the Pacific side reproduced more sexually (c around 50%). Allelic diversity was twice as high in Kamchatka and Kuril populations, suggesting North Pacific coasts being the main last glacial refugia of *A. stella*. We found a stepping-stone pattern of genetic structure from Kamchatka to Atlantic populations, consistent with contemporary ocean currents and melted summer sea ice. Only a subset of the juvenile genetic diversity, mostly of local origin, was found in the established adults, while juveniles exhibited lower levels of genetic differentiation across the Arctic Ocean. Our findings underscore the need for further ecological and behavioral investigations to elucidate the mechanisms allowing the current possibilities of dispersal of this species across the Arctic Ocean.

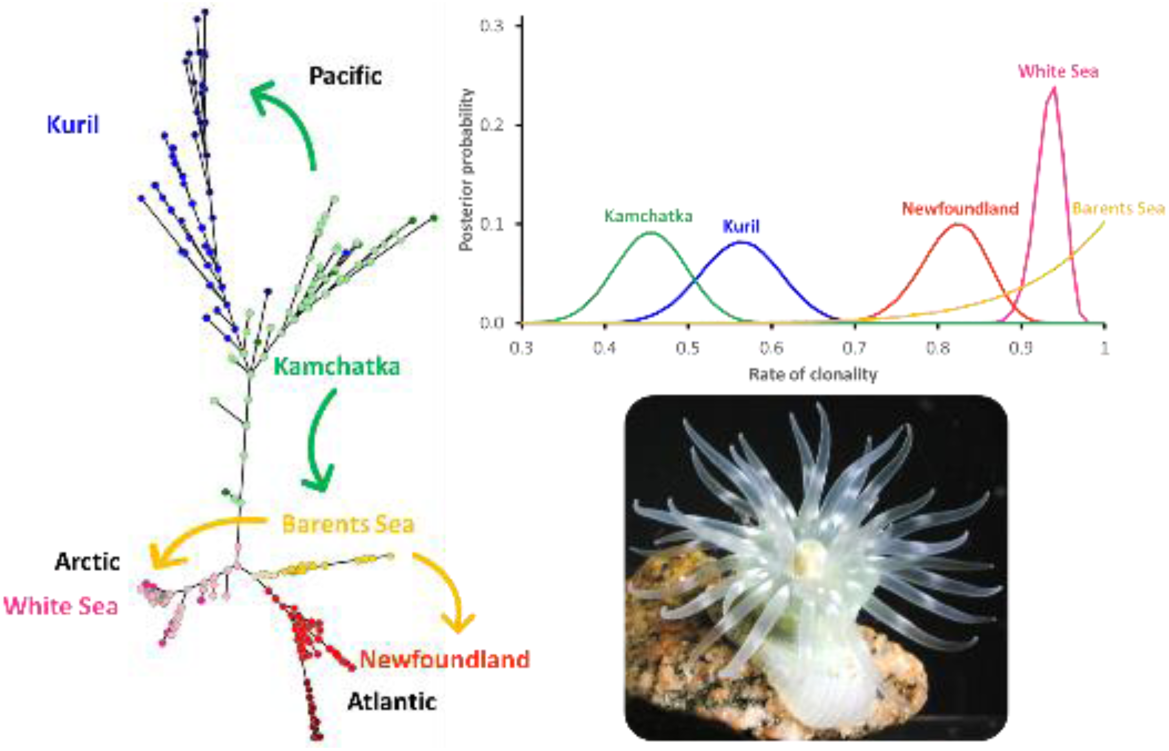

Graphical Abstract (photo: Ekaterina Bocharova)

## Introduction

The way individuals reproduce is the most influential trait in governing genetic diversity and potentially evolutionary response to environmental changes (Duminil et al., 2007; Pierre et al., 2022). Reproductive mode governs the persistence of DNA variations through generations. It has therefore major impacts on the level of genome-wide genetic diversity and its structuration within and among populations (Ellegren & Galtier 2016). Reproductive modes also profoundly affect other biological traits that play important roles for ecological processes, including population growth, dispersal, colonization of new areas and the genetic basis for adaptation (Yu et al., 2016; He et al., 2024).

Clonality is a dominant reproductive mode found in populations colonizing new area, especially when enduring harsh conditions at the edge of their living range (Cadotte et al., 2006; Burke & Bonduriansky, 2018; Roiloa, 2019; Franklin et al., 2021) and in recently invaded, resource-poor environments to which individuals are poorly adapted (geographic parthenogenesis; Peck et al., 1998). All groups and kingdoms of Eukaryota domain, including most of the species that are essential for ecosystem functions, have species that reproduce using partial clonality (Avise, 2015; Schön et al., 2009; Tibayrenc et al., 2015; Orive & Krueger-Hadfield, 2021). In this reproductive mode, parents can generate offspring that are genetically identical to themselves at all loci on their genomes, with the exception of rare recombination events and occasional somatic mutations (De Meeûs et al., 2007; Stoeckel et al., 2021a). In such species, the proportion of descendants produced by clonality, i.e. the rate of clonality, is a key biological factor to understand their genetic and trait evolution (Yu et al., 2016; Becheler et al., 2017). Clonality often increases in populations expanding at range margins, as it allows pioneers to establish and persist in the absence of mates (Baker & Stebbins, 1965; Liu et al., 2006; Wang et al., 2019). Moreover, clonality enables locally successful genotypes to persist across generations without recombining with less adapted ones, thereby accelerating population growth (Fouqueau & Roze, 2021; Pierre et al., 2022). It also decreases the loss of allele diversity and of heterozygosity by genetic drift, especially in small founding populations (Navascues et al., 2010; Stoeckel & Masson, 2014; Reichel et al., 2016; Stoeckel et al., 2021b). However, the absence of recombination in clonal reproduction constrains selection by preventing the combination of adaptive alleles and hindering the removal of deleterious variants (de Visser & Elena, 2007; López-Villavicencio et al., 2013; Pierre et al. 2022).

The Arctic Ocean has been previously considered as an insurmountable barrier for sessile and semi-sessile marine species dispersal due to limited access, harsh environmental conditions and inadequate food resources (Chan et al., 2019). Climate change is opening new routes across the Arctic Ocean, linking northern Pacific and Atlantic biomes (Vermeij & Roopnarine, 2008). The recent acceleration of summer sea-ice loss, now surpassing levels recorded since the Holocene Climatic Optimum (∼8,000 years before present, estimated at +0.3 °C relative warming above pre-industrial levels; compared to +1.34 °C in the Arctic in 2024; Laakkonen et al., 2021; E.U. Copernicus Global Climate Highlights, 2024), is likely to facilitate renewed secondary contact between species and populations from the northern Atlantic and Pacific biomes (Brandt et al., 2023). Understanding genetic diversity and population structure in such species provide useful insights into the ecological and evolutionary processes that occur during range expansion under real and complex conditions (Sax et al., 2007; Moran & Alexender, 2014; Bridle & Hoffmann, 2022). It also provides a frame of reference to managers and policymakers to identify priorities and mechanisms for mitigating adverse effects of global changes on intraspecific diversity and for controlling future invasive populations (Chan et al., 2019).

Sea anemones are semi-sessile organisms reproducing using partial clonality (Bocharova & Kozevich, 2011; Bocharova, 2016). *Aulactinia stella* (Cnidaria, Anthozoa) is an intertidal and sublittoral species belonging to Actiniidae family. It was first observed and described on the eastern coast of the USA in the 19th century (Verrill, 1864), probably in the southernmost part of its distribution area. Its geographic range is rather circumpolar, along the coasts of the Arctic Ocean and of the northern parts of Atlantic and Pacific Oceans (Grebelnyi, 1980; Sanamyan & Sanamyan, 1998, 2006). First studies of *A. stella* populations in the Arctic Ocean, on the Murman Coast, were carried on about fifty years ago (Loseva, 1974). Adults of *A. stella* mainly live fixed to rocky ground, even if it can occasionally be found buried in sandy or muddy soils, hidden in crevices or fixed to debris (Bocharova, 2012). It feed predating smaller mollusks, crustaceans, fish larvae and other animals (Bocharova, 2012). The population density of *A. stella* ranges from 1 to 18 individuals per square meter in the littoral zone of the White Sea, and from 1 to 14 individuals per square meter along the Barents Sea coasts (Bocharova, 2012). In laboratory conditions, adults can survive for several decades, but their natural lifespan in the wild still remains unknown.

The life cycle of *A. stella* includes a swimming larval stage, which can be brooded by the parental anemone until it develops into a polyp. The polyp stage is benthic, but it retains the ability to be transported passively by water currents until it settles as adult on a solid substrate. In response to unfavorable abiotic or biotic conditions, many sea anemone species are capable of detaching from the substrate and temporarily drifting with tidal and coastal currents, a behavior that is particularly common when winter conditions become extreme (Riemann-Zürneck, 2008). Some circum-Antarctic and Arctic sea anemones present a distinct post-metamorphic juvenile stage, following the larval phase, during which they drift within pelagic currents (Riemann-Zürneck, 2008). To date, no comparable pelagic stage has been documented or investigated for *A. stella* despite the presence of a free-living polyp stage within the benthic fauna with similar potential (Bocharova, 2012). Reproduction occurs annually from May to October, with phenological variation depending on local environmental conditions, but a peak period between June and August (Bocharova, 2012). Along the Kamchatka coasts during this time, males release gametes passively into the water column, that are later ingested by females. Females brood the developing larvae internally (Bocharova, 2012). As for many circumpolar marine species, direct observations of reproductive and dispersal processes still remain unrealistic. Consequently, genetic analyses remain the most effective approach for inferring key biological and ecological traits of this species in natural conditions.

Developmental and histological evidence suggests that this species may reproduce through parthenogenesis (Bocharova, 2012). Preliminary analyses based on a limited number of mitochondrial and ribosomal sequences hypothesized that these populations are likely highly clonal and exhibit strong genetic structuring across the Arctic Ocean despite the region’s increasing ice melt over recent decades (Bocharova & Mugue, 2012; Bocharova, 2015). While these sequences were suitable for phylogenetic analyses, they lacked the resolution necessary required to confirm clonality and accurately characterize the genetic structure of *A. stella* populations. The recent development of microsatellite primer pairs now enables investigating the biology of this circumpolar sea anemone more precisely, using approaches based on genetic diversity and its spatial patterns (Bocharova et al., 2018).

In this study, we aimed to confirm the occurrence of clonal reproduction in natural populations of *A. stella*, quantify its extent and variability across the species’ range, and use high-resolution, polymorphic markers to accurately assess population structure throughout the Arctic Ocean. To investigate clonal reproduction in this species, we analyzed genotypic and genetic indices across five geographically distinct sites spanning the Arctic Ocean, including locations in the northern Atlantic and northern Pacific. We leveraged previous sampling campaigns (2012–2017), which included fewer sites and greater temporal gaps in the Pacific. While this may influence some interpretations, our results still provide a robust overview of the species’ reproductive variation and current biogeography, especially given its laboratory lifespan of at least more than one decade.

A total of 314 individuals, representing two age cohorts, juveniles and adults, were sampled to estimate the contribution of clonal reproduction and its variation across the species’ range. By examining temporal changes in genotypic frequencies within each site, we aimed to infer the prevalence of clonal reproduction. We hypothesized that the limited energetic and physiological cost of clonal reproduction may facilitate the establishment and persistence of populations under harsh polar coastal conditions (Peck et al., 1998; Klimeš, 2008; Wang et al., 2018). Conversely, we also considered that the genetic recombination associated with sexual reproduction may promote the emergence of genotypes better adapted to local environments (Pierre et al., 2022). Our findings on reproductive modes are interpreted in the context of the specific ecological challenges faced by species expanding into previously unfavorable habitats. In addition, we used the genotyped populations to assess genetic differentiation and identity in state between individuals and sampled sites. Sampling across the full distribution range allowed us to pinpoint regions with higher genetic diversity, which may correspond to historical refugia. By comparing genetic differentiation between juveniles and adults, we aimed to identify present-day or historical corresponding dispersal routes. Although *A. stella* was first described along the North Atlantic coasts, suggesting a possible Atlantic origin, previous limited sequence data instead indicated higher genetic diversity in North Pacific populations. If confirmed using more polymorphic and resolutive markers that are microsatellites, this may suggest a more recent expansion into the North Atlantic via the Arctic Ocean. Considering the free-living polyp stage which can disperse passively via marine currents, as well as the potential for semi-sessile adults to be transported on drifting debris or through human-mediated movement of materials, we expect a limited but ongoing level of gene flow facilitated by favorable summer sea conditions. Under this scenario, we would expect to observe shared probabilities of identity in state among populations and a stepping-stone pattern of genetic differentiation across the Arctic, potentially with preferential dispersal routes connecting sampled populations. Alternatively, considering populations evolving isolated due to current Arctic conditions for ∼8,000 years (Holocene Thermal Maximum) or even ∼125,000 years (last interglacial Eemian period), we would expect very distinct and strongly structured genetic diversity with even fixed and private alleles between populations (Slatkin & Takahata, 1985), out of artefactual homoplasy.

Ultimately, our findings will provide a baseline framework for understanding the genetic structure of this species across its Arctic distribution, offering critical insights at a time of profound environmental change in this region (Laakkonen et al., 2021; Brandt et al., 2023).

## Materials and methods

### Sample collection

In September 2012, we sampled at the low tide in the littoral zone 23 *Aulactinia stella* in the White Sea Biological Station of Moscow State University (WSBS), Kandalaksha Bay of the White Sea and 30 individuals at Liinkhamari, in the Varanger Fjord of the Barents Sea (see Bocharova 2015). We also sampled using SCUBA diving at 5-18 m depth in the sublittoral zone 29 individuals at the Cape Broyle in the Admirals Cove area, Newfoundland, Canada, in June 2017. Along Kamchatka coast of the Pacific Ocean, we collected 5 individuals in the Avachinsky Bay in June-July 2014 and 9 individuals in the Viluchinsky Bay in June-July 2016 at 5-21 m depth. We used these 14 Kamchatka adult individuals for mitochondrial DNA sequencing, but only 6 of them brooded juveniles. Therefore, for microsatellite analysis, we additionally analysed 3 females with their brooded offspring collected in the Avachinsky Bay in July 2012 (2 samples) and June 2013 (1 sample). At low tide, we sampled 20 individuals in the intertidal zone of the Matua Island of the Kuril Islands in August 2016 (see map of the sampled sites on Figure S1).

We checked that the phenotypic and biometric appearance of each individual belonged to *A. stella* species using descriptions in Loseva (1974), Fautin Dunn et al. (1980), Sanamyan and Sanamyan (2008), Sanamyan et al. (2019, 2020) and WoRMS (Rodríguez et al., 2025). In addition, we used nematocyst identification from tentacles, pharynx, body wall and mesentery (Manuel, 1981), and mitochondrial DNA sequence analysis to confirm the species. To analyze genetic diversity in two successive generations, we purposely genotyped large females and the juveniles they brooded in their gastrovascular cavity. The quantity of juveniles in such females varied from 1 to 39 per female and we sequenced them all.

### DNA extraction, mtDNA sequencing and genotyping by microsatellite loci

We extracted the total DNA of each sampled adult and juvenile polyp. For each adult, we used a part of the adult pedal disk measuring approximately 3x3x3 mm; And for juveniles, we extracted DNA from the whole juvenile polyps which measured from 1 to 5 mm. For each sample, we desiccated and grinded the sampled tissues. We then used the Wizard SV Genomic DNA Purification System (Promega, USA) following the manufacturer’s protocol. Extracted DNA was preserved in 96% ethanol before genotyping.

For mitochondrial gene (12S rRNA, 16S rRNA, cytochrome oxidase III) sequencing, we only used adult subsamples: 23 individuals from the White Sea, 30 from the Barents Sea, 29 from Newfoundland, 14 from Kamchatka, and 20 from the Kuril Islands. We applied the primer sets and methodology thoroughly described for 16S rRNA and COIII (Geller & Walton, 2001) and 12S rRNA mitochondrial fragments (Bocharova 2015). Sequencing was performed by 3500 Genetic Analyzer (Applied Biosystems, USA) using BigDye Terminator v3.1 Cycle Sequencing Kit. The sequences were analyzed by Geneious Prime software. By concatenating these three mtDNA sequences (12S, 16S, COIII), we grouped the samples into haplotypes that were aligned and perfectly matched the five previously described haplotypes (Bocharova, 2015). Including these mitochondrial haplotypes (Pacific 1-5), we used geneHapR (Zhang et al., 2023) package to produce the haplotype network (Figure 2). We then used the extracted nuclear DNA to amplify 41 microsatellites using primer pairs previously developed in Bocharova et. al. (2018) for this species and another sea anemone species of Actiniidae family, *Cribrinopsis albopunctata* (Sanamyan & Sanamyan 2006, Sanamyan et al. 2019) with the corresponding PCR conditions. PCR products were visualized in 3500 Genetic Analyzer (Applied Biosystems, USA) using POP7 gel polymer. We analyzed the obtained chromatograms using GenMapper Software (ThermoFisher Scientific, USA). Out of these 41 primer pairs, only ten pairs (Act007, Act011, Act028, Act067, Act173, Act177, Act235, Act238, Act249, Act386,) successfully amplified and were polymorphic in our samples. We then multiplexed the ten microsatellites into four PCR pools using three specific dyes (TAM, ROX, FAM) for visualization: Pool 1) Act007 (FAM), Act011 (TAM); Pool 2) Act238 (FAM), Act249 (TAM), Act173 (ROX); Pool 3) Act067 (FAM), Act177 (ROX), Act386 (TAM); Pool 4) Act28 (TAM), Act235 (ROX).

To estimate if this marker set was powerful enough to distinguish between individuals, we computed the overall observed unbiased probability of identity in state under panmixia (hereafter overall pidu) and the overall observed unbiased probability of identity in state between sibs (overall pids) as recommended by Waits et al. (2001). To achieve allele dosage within each individual per locus, we used the dose-effect method with some modifications (Rodzen & Mаy, 2002; Drauch Schreier et аl., 2011). In cases of more than 1 allele, dose-effect is counted following the instruction (Figure S2): two alleles of almost the same height were considered as AABB (Figure S2A); One high allele and two low alleles as AABC (Figure S2B); One very high allele and one low allele as AAAB (Figure S2C); Tree alleles of the same height were considered as ABC + null allele (Figure S2D); One high allele and one allele half lower were considered as AAB + null allele (Figure S2E).

### Analyses of genetic diversity within spatio-temporal samples

Parents that produce descendants by clonality can result to observe multiple times the same Multi-Locus Genotype (hereafter MLG) among the genotyped samples (Halkett et al., 2005; Arnaud-Haond et al., 2007). However, these repeated MLGs may also come from genetic marker lacking sufficient resolution to disentangle between descendants produced by sexual reproduction, either under panmixia (the lower probability to observe two identical sexual descendants) or between mating sibs (the higher probability to observe two identical sexual descendants). To estimate the power of our marker set to distinguish different individuals and assess the upper and lower bounds for the probability of observing identical MLG drawing a pair of individuals sampled in a same population, we therefore computed the probabilities of identity under panmixia (*pidu*) and considering only sib mating (*pids*; Jacquard, 1968, 1970; Waits et al., 2001) using formula adapted for tetraploid populations implemented in GenAPoPop software (Stoeckel et al., 2024).

To assess the level of genetic diversity within populations and compare between populations, for each age cohort in one site, we reported the sample size (*N*) and computed the average number of observed alleles 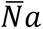 and its variance Var(Na) over the ten microsatellites, the average effective number of alleles 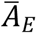 and its variance Var(*A*_*E*_), the average gene diversity 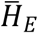 and its variance Var(*H*_*E*_), the average observed heterozygosity 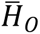 and its variance Var(*H*_*0*_) and the average value of the inbreeding coefficient 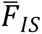 and its variance Var(*F*_*IS*_) over all loci.

We also computed genotypic indices including the number of unique genotypes (G), the unbiased genotypic richness R (Dorken & Eckert, 2001), the genotypic evenness as Pareto β to assess the potential uneven contribution of some clonal MLGs to the genotypic composition of populations (Arnaud-Haond et al., 2007; Bailleul et al., 2016).

We searched for clonal signatures in genetic diversity by analyzing 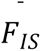 and its variance Var(*F_IS_*) and the overall estimate of the linkage disequilibrium (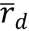, Agapow & Burt, 2001) as commonly interpreted using one-shoot samples (Halket et al., 2005; Arnaud-Haond et al., 2020; Stoeckel et al., 2021a). Finally, to assess rates of clonality within spatial sites, we used the Bayesian estimate of rates of clonality using genotype frequency changes between adult and juvenile at the same sample site (ClonEstiMate, Becheler et al., 2017 adapted to autpolyploids in GenAPoPop Stoeckel et al., 2024).

To assess the genetic structure between populations and generations, we computed temporal and spatial *pST* (Ronfort et al., 1998) using GenAPoPop, and plotted the unrooted minimum spanning tree between individuals using their pairwise number of shared alleles (identity-in-state) using the classical equal-angle algorithm (Felsenstein, 2004) to picture the coancestry between individuals across populations at our marker set. To assess the dynamics (speed and direction) of genetic diversity along sampling sites, we compared juvenile and adult subpopulations sampled at the same sites and between sites computing the corresponding *pST*. We also used a Discriminant Analysis of Principal Components (DAPC, Jombart et al., 2010) to investigate the genetic structure of our five sites and tens cohorts. We chose to retain 80 principal component eigenvalues. We used a knee locator approach to find the minimal number of clusters best structuring the differentiation between individuals and groups. We verified this value by increasing and decreasing the number of clusters and checking that we obtained the same groups and distribution of clusters among sites and cohorts.

### Statistical analyses

To test if distribution of population genetic indices were significantly different between Arctic-Atlantic and Pacific populations and between juveniles and adults, we used Mann-Whitney test and reported the median values of the two groups (*median_group1_* versus *median_group2_*) in the order they are mentioned, the sum of the rank of the first group (*U*), the number of measures in each group (*n1=4* for the cohorts sampled on the Pacific side of the Arctic Ocean and *n2=6* for the cohorts sampled on the Atlantic side) and the probability value (*p*) of the hypothesis that the two groups have identical distribution of their genetic indices. To compare *pST* between juveniles and adults among pairs of the same sites, we used Wilcoxon signed rank test. We reported the median *pST* value of the two groups (*median_group1_* versus *median_group2_*) in the order they are mentioned, the sum of the ranks of the differences above zero (*W*) and the probability value (*p*) of the hypothesis that the two age cohorts have identical distribution of *pST*. All these tests were performed using python 3.11 and Scipy.stats 1.13 (Virtanen et al., 2020).

## Results

We genotyped 314 individuals of *Aulactinia stella* sampled on five sites, including a total of 75 adults, ranging from 9 to 21 with a median of 15 adults per site, and a total of 239 juveniles, ranging from 6 to 93 with a median of 49 juveniles per site (Table 1). We found both females and males in the Kamchatka population, whereas we found only females in the Kuril Islands, Newfoundland, the Barents and the White Seas. For subsequent genetic analyses, we only used the adult females from each site.

**Table 1:** Summary of population genetic indices in ten spatio-temporal populations of *Aulactinia stella*. For each age cohort at one site, are reported the sample size *N*, number of genotypes *G*, genotypic richness *R*, genotypic evenness *Paretoβ*, overall estimate of linkage disequilibrium 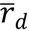, average number of observed alleles *Na* and its variance *V(Na)* over the ten microsatellites, average effective number of alleles 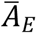 and its variance V(*A*_*E*_), average observed heterozygosity 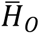 and its variance V(*H*_*0*_), average gene diversity 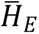 and its variance V(*H*_*E*_), average value of the inbreeding coefficient 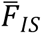 and its variance V(*F_IS_*) over all loci, unbiased probability of identity under panmixia *pidu* and between sibs *pids*.

| Site | Age cohort | N | G | R | Paretoβ | $\bar{r}_d$ | Na | V(Na) | $\bar{A}_E$ | V($A_E$) | $\bar{H}_O$ | V($H_O$) | $\bar{H}_E$ | V($H_E$) | $\bar{F}_{IS}$ | V($F_{IS}$) | pidu | pids |
| --- | --- | --- | --- | --- | --- | --- | --- | --- | --- | --- | --- | --- | --- | --- | --- | --- | --- | --- |
| Kamchatka | Adults | 6 | 6 | 1.000 | 1.807 | 0.011 | 3.0 | 30 | 2.2 | 17.27 | 0.294 | 0.488 | 0.416 | 0.590 | 0.292 | 1.635 | 1.1E-8 | 0.0043 |
|  | Juveniles | 49 | 48 | 0.979 | 4.615 | 0.122 | 5.2 | 151.6 | 2.4 | 24.25 | 0.314 | 0.397 | 0.460 | 0.534 | 0.318 | 1.156 | 1.6E-7 | 0.0093 |
| Kuril Islands | Adults | 19 | 19 | 1.000 | 3.322 | 0.026 | 4.6 | 84.4 | 2.5 | 46.90 | 0.176 | 0.121 | 0.438 | 0.640 | 0.598 | 1.016 | 9.9E-9 | 0.0040 |
|  | Juveniles | 30 | 30 | 1.000 | 3.954 | 0.077 | 5.7 | 234.1 | 2.5 | 53.09 | 0.188 | 0.301 | 0.372 | 0.891 | 0.494 | 0.948 | 1.9E-9 | 0.0026 |
| Barents Sea | Adults | 21 | 10 | 0.450 | 0.302 | 0.444 | 2.5 | 14.5 | 1.4 | 0.98 | 0.197 | 0.425 | 0.223 | 0.287 | 0.115 | 3.124 | 3.8E-4 | 0.1901 |
|  | Juveniles | 6 | 2 | 0.200 | 0.591 | 1.000 | 1.2 | 1.6 | 1.2 | 1.42 | 0.125 | 0.628 | 0.097 | 0.376 | -0.290 | 13.323 | 0.0776 | 0.5125 |
| Newfoundland | Adults | 14 | 12 | 0.846 | 1.807 | 0.025 | 2.1 | 18.9 | 1.4 | 6.47 | 0.165 | 0.656 | 0.170 | 0.512 | 0.024 | 4.971 | 0.0011 | 0.1556 |
|  | Juveniles | 61 | 28 | 0.450 | 0.716 | 0.076 | 1.8 | 9.6 | 1.4 | 6.63 | 0.158 | 0.747 | 0.178 | 0.596 | 0.112 | 4.875 | 0.0019 | 0.1606 |
| White Sea | Adults | 15 | 5 | 0.286 | 0.129 | 0.619 | 1.9 | 18.9 | 1.6 | 7.81 | 0.314 | 1.526 | 0.238 | 0.886 | -0.322 | 10.807 | 2.3 E-4 | 0.0635 |
|  | Juveniles | 93 | 28 | 0.293 | 0.261 | 0.268 | 3.0 | 48 | 1.7 | 8.53 | 0.308 | 1.447 | 0.266 | 0.912 | -0.157 | 7.841 | 3.9E-5 | 0.0358 |
| Over All | Both | 314 | 180 | 0.572 | 0.482 | 0.293 | 11.1 | 468.9 | 2.1 | 6.24 | 0.243 | 0.387 | 0.440 | 0.454 | 0.448 | 0.870 | 8.3E-9 | 0.0045 |

We found 15 brooding females in White Sea, 21 in Barents Sea, 14 in Newfoundland, 9 in Kamchatka and 19 in Kuril Islands sampled sites. The oral disc diameter of mature females varied from 10 to 45 mm while the overall size of the juveniles released from the gastrovascular cavities of the adults varied from 1 to 5 mm.

### Gene, allele and genotypic diversities in nuclear microsatellites

Gene, allele and genotypic diversities in nuclear microsatellites were consistent over generations (Table 1). We found a maximum of four different alleles for each locus in one individual over all our samples and concluded that *A. stella* must be at least an autotetraploid species. Using our ten microsatellites, the mean numbers of alleles per locus per site and developmental stage ranged from 1.2 to 5.7, with a median of 2.75. The mean effective numbers of alleles per locus per site and developmental stage ranged from 1.19 to 2.53 with a median of 1.69. Over all samples, probability of identities under panmixia was 8.3×10^-9^ and probability of identity between sibs was 4.5×10^-3^. These low probabilities that two individuals generated by sexual reproduction would result into similar genotypes advocated that the ten microsatellites we used were powerful enough to genetically distinguish between sexual descendants, even resulting from a sib mating (Waits et al. 2001). We thus considered afterward that individuals with identical MLGs were individuals of the same clone (i.e., ramets of a same genet).

Over all sites and cohorts, we found 154 unique MLGs and 26 of these MLGs were found repeated in more than one sampled individual, i.e. genets with more than one ramet (Table S1). In Kuril Islands, we genotyped no repeated MLG. In Kamchatka, we found three individuals of the same clone. In Barents Sea, we found three repeated MLGs, including respectively two, five and thirteen ramets. In White Sea, we found eight repeated MLGs involving two to forty ramets, for a total of 88 ramets. In Newfoundland, we found fourteen repeated MLGs involving two to eleven ramets, for a total of 49 ramets. All ramets of repeated MLGs were exclusively sampled on the same site. In all populations with repeated MLGs, repeated MLGs were found both in adults and in juveniles. Some of these genets were even shared by adults and juveniles, except in the Newfoundland samples.

We found higher genetic and genotypic diversity, and lower probabilities of identity on the Pacific side sites (i.e., Kuril and Kamchatka) than found on the Atlantic side of the Arctic Ocean (i.e., Newfoundland, Barents and White Seas), even if differences among sites within oceans still remained on some indices (Table 1). The four cohorts sampled in Kuril Islands and in Kamchatka showed nearly twice more average gene diversity, ranging from 0.372 to 0.460, than found in the six cohorts sampled in Newfoundland, Barents and White Seas, that ranged from 0.097 to 0.266 (0.427 versus 0.174, U=24, n1=4, n2=6, p=0.009), while they both showed similar standard deviation of gene diversity over all loci (U=15, n1=4, n2=6, p=0.610; Table 1). Cohorts in the Pacific sites also showed significantly higher mean numbers of alleles per locus (4.9 versus 2.0, U=23.5, n1=4, n2=6, p=0.019) and higher effective number of alleles (2.43 versus 1.43, U=24, n1=4, n2=6, p=0.009).

Kamchatka and Kuril samples showed higher genotypic richness measured as R than populations on the Atlantic side (1.00 versus 0.37, U=24, n1=4, n2=6, p=0.009) and higher genotypic evenness measured as Pareto β values (3.64 versus 0.45, U=23.5, n1=4, n2=6, p=0.019). Over all loci, Kamchatka and Kuril samples showed lower probabilities of identity under panmixia (<10^-6^ versus 0.003, U=0, n1=4, n2=6, p=0.009) and between sibs (0.008 versus 0.118, U=0, n1=4, n2=6, p=0.009) than found in samples from the Atlantic side.

### Heterozygosity and deviation from the genotypic proportions expected under Hardy-Weinberg hypotheses

We found no difference in observed heterozygosity between cohorts on the Pacific and Atlantic sides (0.241 versus 0.181, U=15, n1=4, n2=6, p=0.61), though variance of observed heterozygosity between loci was nearly two times higher in cohorts on the Atlantic side (0.389 versus 0.701, U=1, n1=4, n2=6, p=0.019). Mean Fis over all loci were significantly lower in Newfoundland, the Barents and the White Seas than in Kamchatka and Kurils (0.406 versus - 0.066, U=24, n1=4, n2=6, p=0.009) and even negative in the juvenile cohort of Barents Sea, and in adults and juveniles of White Sea. Variances of Fis values between loci were significantly higher, three to ten times, in cohorts on the Atlantic side (1.08 versus 6.41, U=24, n1=4, n2=6, p=0.009), than found in populations on the Pacific side. Though linkage disequilibrium seemed higher in cohorts on the Atlantic side, they were not significantly different from values measured in cohorts on the Pacific side (0.051versus 0.356, U=5, n1=4, n2=6, p=0.171).

### Estimated rates of clonality using temporal genotype frequencies

Credible intervals including 99% of posterior probabilities of rates of clonality estimated from temporal samples were quite sharp, spanning around their peaks on a range of 0.06 to maximum 0.25 (Figure 1). Clonality was intermediate in the Kuril Islands and Kamchatka, but very high in the Barents Sea, the White Sea, and Newfoundland.

**Figure 1:**
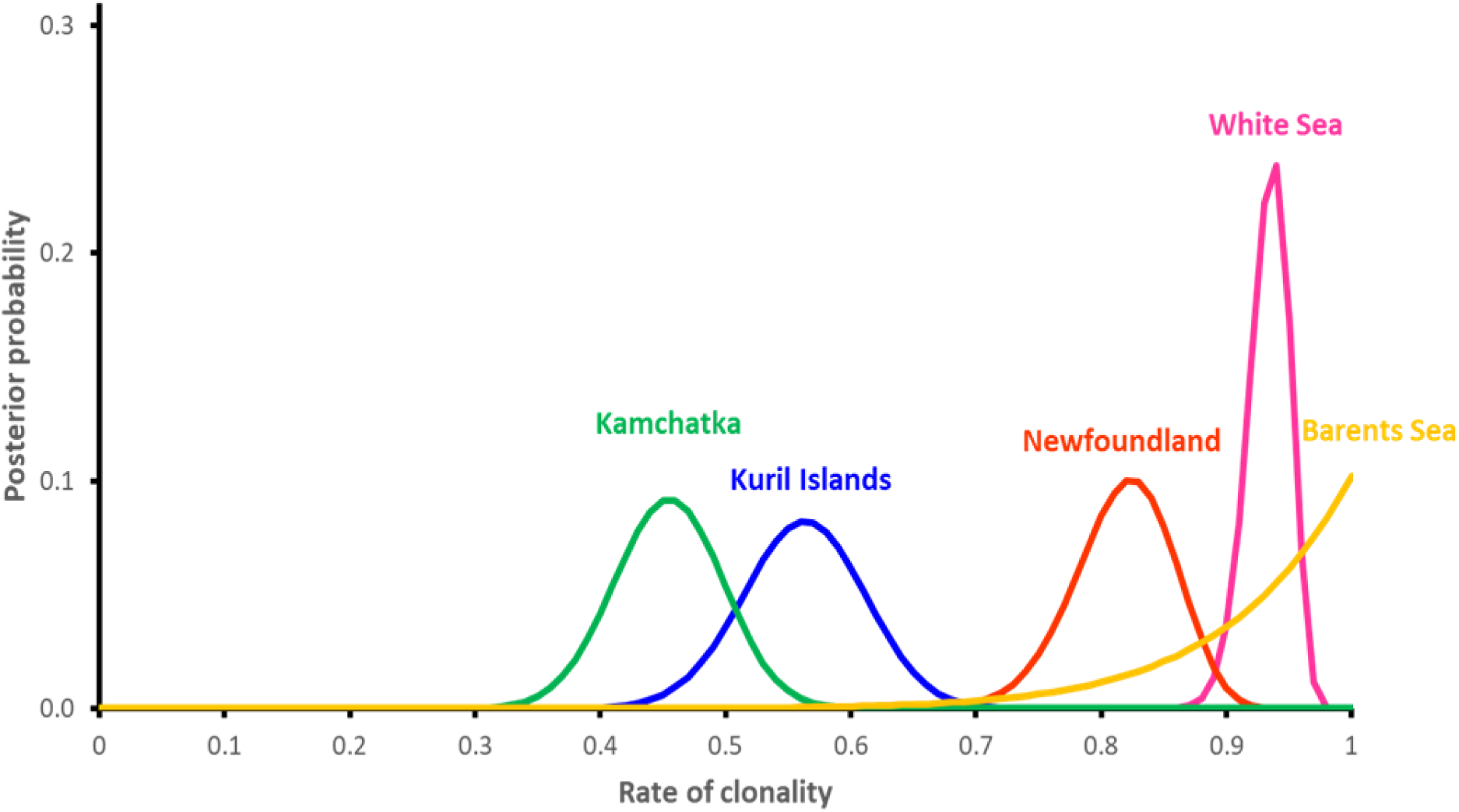
Posterior density probabilities of estimated rates of clonality using changes of genotype frequencies over one generation for each sampled site using ClonEstiMate method for autopolyploids in GenAPoPop software (Becheler et al., 2007; Stoeckel et al., 2024) on the ten nuclear microsatellites. Kuril Islands in blue, Kamchatka in green, Newfoundland in red, White Sea in pink and Barents Sea in gold.

### Genetic structure between sites using mitochondrial gene sequences

Using three gene sequences (12S, 16S, COIII), we found that all the 82 adults sampled on the Atlantic side (Newfoundland, the White and the Barents Seas) shared the same mitochondrial haplotype, Pacific-3 (classification according to Bocharova, 2015; GenBank Accession Numbers: 12S: KT310190, 16S: KT310200, COIII: KT310212). On the Pacific side, all the 20 individuals sampled in the Kuril Islands belonged to another mitochondrial haplotype, Pacific-1 (GenBank Accession Numbers: 12S: KT310188, 16S: KT310198, COIII: KT310210) while Kamchatka’s samples distributed among five mitochondrial haplotypes (Figure 2).

**Figure 2:**
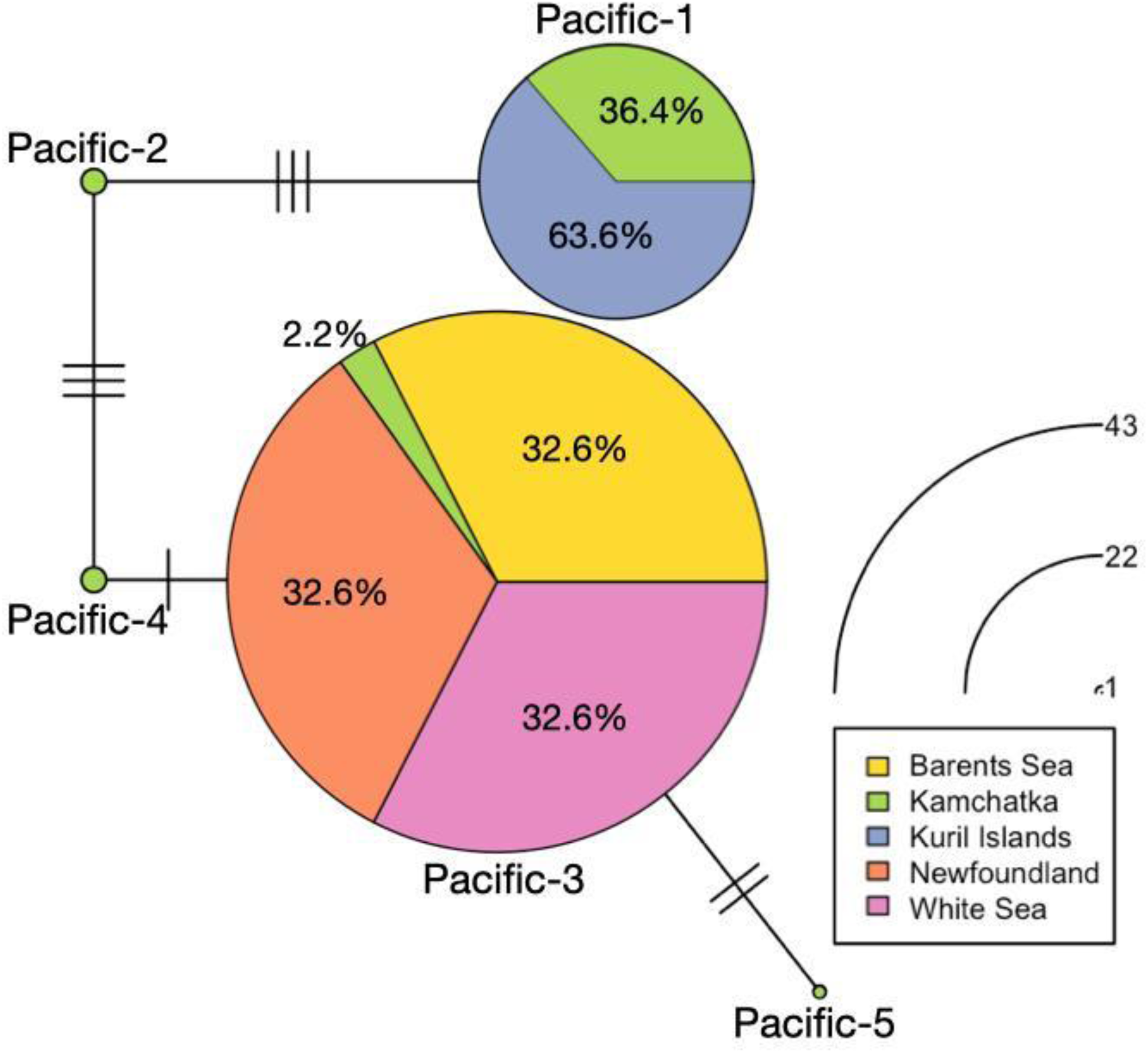
Haplotype network using three mitochondrial gene sequences (12S, 16S, COIII). The proportion of each sampled site per haplotype is represented in color. The circle sizes of each haplotype are proportional to their number in our dataset. We used haplotype names initially identified in Bocharova (2015), thus Pacific 1 to 5. We found no new haplotypes.

### Genetic differentiation between sites and cohorts using nuclear microsatellites

Among adults, genetic differentiation between sites ranged from 0.155 to 0.612 while among juveniles, genetic differentiation between sites ranged from 0.105 to 0.530 (Table S2). Within each sampled site, *pST* between adults and juveniles ranged from 0.009 to 0.088, therefore between six to eleven times lower than between sites. Between sites, adults were more genetically structured than juveniles (0.366 versus 0.245, W=43, p=0.065; Figures 3, S1, S3). Considering the adults and juveniles together by site, genetic differentiation between sites ranged from 0.096 to 0.521 (Table S2a). Globally, in juveniles, *pST* between sites showed very close values and followed ranks of geographical and along-the-coast distances (Table S2b, Figure S1). In contrast, adults of White Sea appeared highly genetically isolated from the other sites, excepted from juveniles in Newfoundland. *pST* genetic differentiation hierarchically structured the samples into one Pacific side group and one Atlantic side group then subdivided by sites and then by age cohorts (Table S2b, Figure 3).

**Figure 3:**
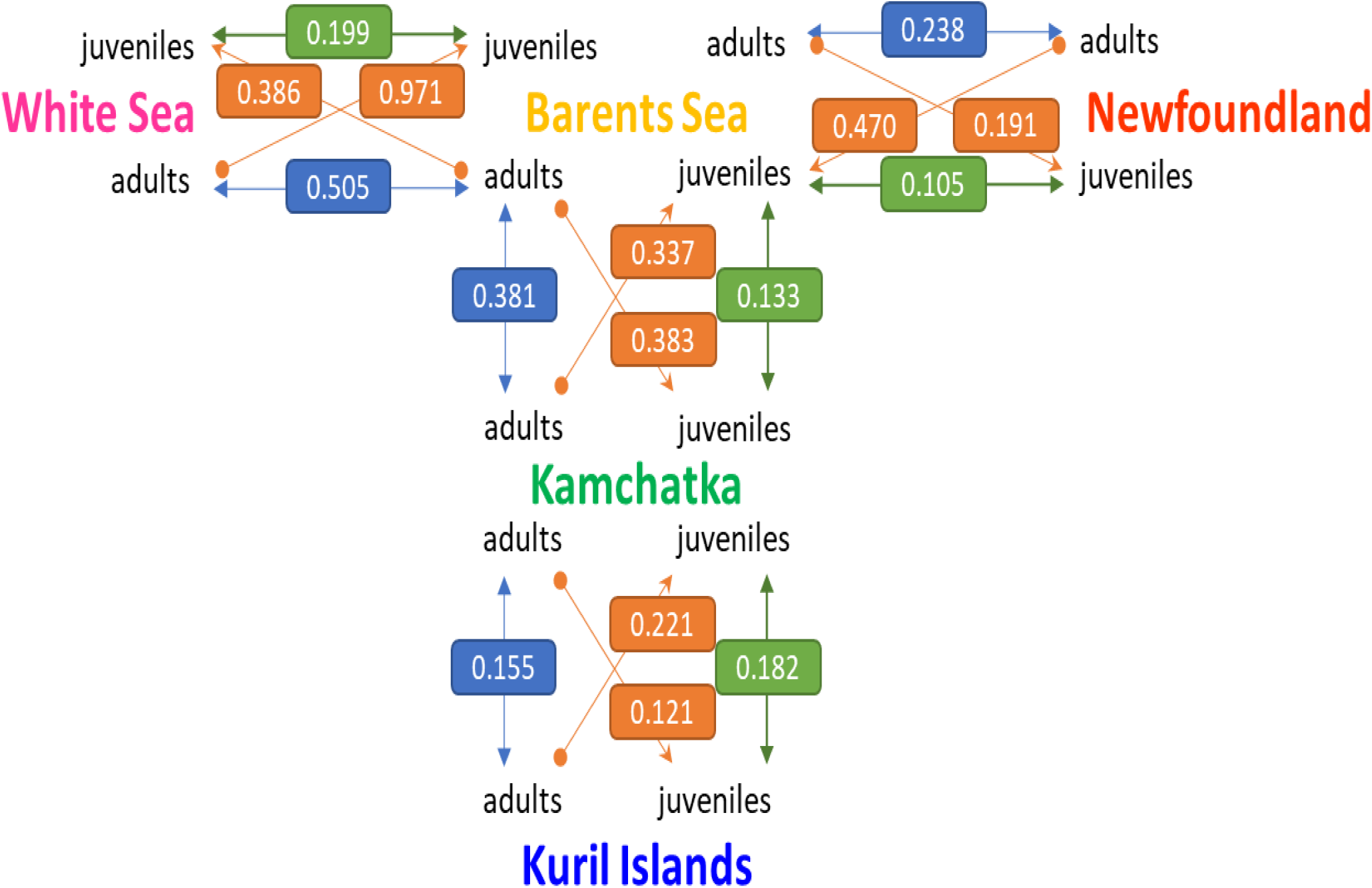
Oriented graph of *pST* values among sites and cohorts (vertices), based on ten nuclear microsatellite loci. Orange arrows (edges) and the *pST* values in boxes illustrate pairwise genetic differentiation between sites, allowing inference of the dynamics and direction of genetic spread. Comparisons of *pST* values among adults (blue boxes) and juveniles (green boxes) between site pairs reveal temporal genetic differentiation, even if apparent increases or decreases in genetic differentiation may arise by chance due to sampling variability within the same underlying distribution, or as a result of homoplasy or gene flow between populations. For instance, adults from the Barents Sea are twice as identical in state with Newfoundland juveniles as adults from Newfoundland are with Barents Sea juveniles, suggesting preferential gene flow from the Barents Sea toward Newfoundland. Moreover, genetic differentiation between Newfoundland and Barents Sea adults is approximately twice that observed between juveniles, indicating either enhanced gene flow among juveniles or processes promoting endemism in established adults.

This hierarchical genetic structure was congruently supported by the DAPC results (Figure 4). Interestingly, the Bayesian information criteria of the DAPC decreased continuously with the increasing number of clusters, as expected in a spatial stepping-stone model (Figure S4). The minimal number of clusters maximizing the distance between groups was 4. Whatever the number of clusters, DAPC grouped together individuals of Barents Sea and Newfoundland into one common and compact group, while the other sites each stood for their own genetic groups with couples to handful individuals assigned to other groups. Looking at the genetic structure between cohorts and sites, juveniles from the Kuril Islands shared more genetic diversity with adults from Kamchatka than the inverse (Figure 2). Adults of Barents Sea shared more similar genetic diversity with juveniles of Newfoundland and of White Sea than the inverse. DAPC proportion of each individual assigned to one DAPC cluster also supported this dynamic tendency (Figure 4): one descendant sampled in Kuril was assigned to Kamchatka genetic cluster; Three adults sampled in Kamchatka were assigned to Barents-Newfoundland genetic cluster and one adult sampled in Barents Sea was assigned to Kamchatka genetic cluster; Four adults sampled in Barents Sea were assigned to white sea genetic cluster, and one adult and ten descendants sampled in White Sea were assigned to Barents-Newfoundland genetic cluster.

**Figure 4:**
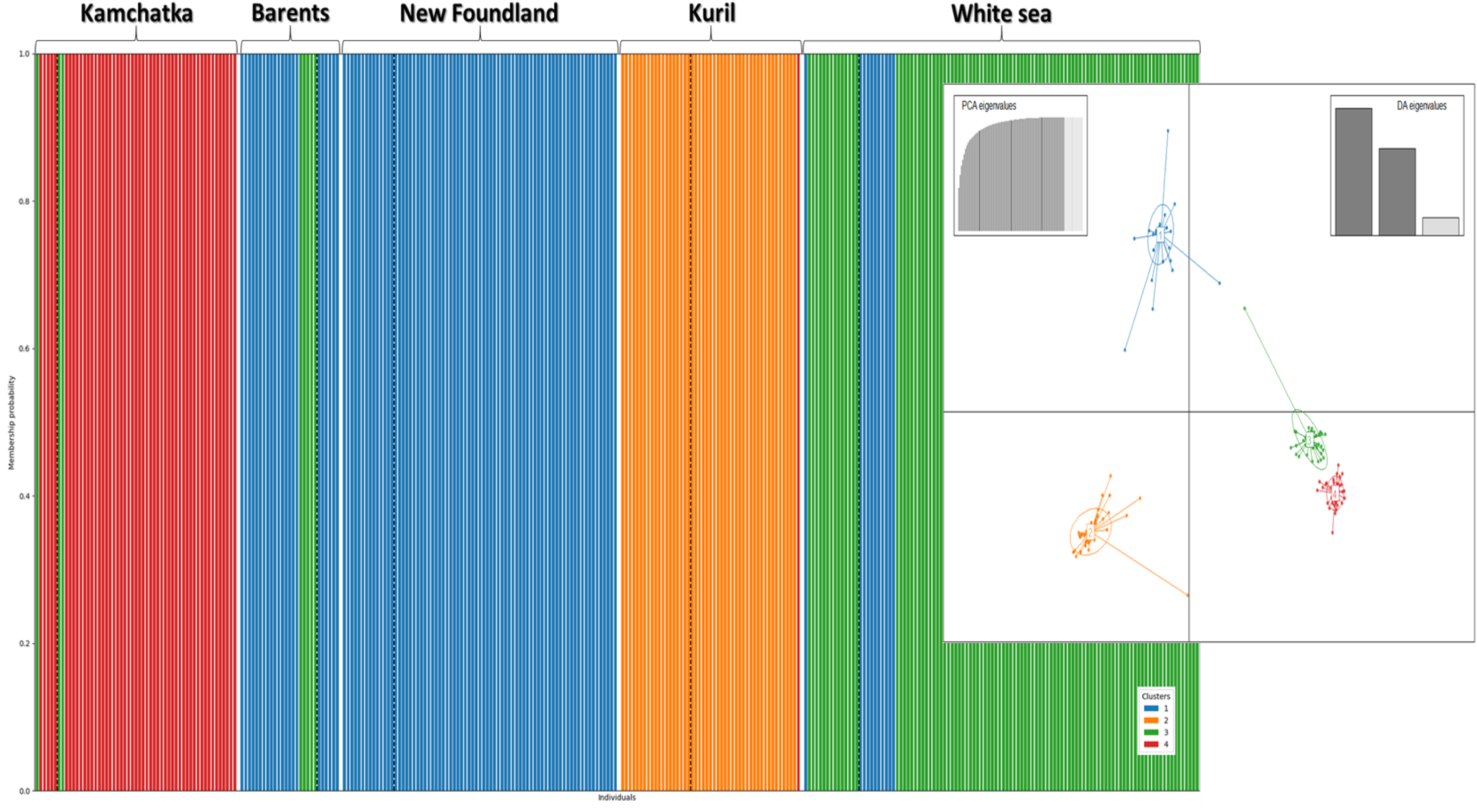
Discriminant Analysis of Principal Components of the genetic structure of *Aulactinia stella* across the Arctic Ocean using the ten nuclear microsatellites. We found four clusters being the minimal number of clusters maximizing the distance between groups (see Figure S4). Individuals are represented as columns corresponding to the membership probability to the four supported clusters (in color). Sampled sites are represented successively, separated by a large white bar. Within each site, adults are on the left side of the dashed vertical black line and juveniles on the right.

Similarly, the minimum spanning tree of pairwise genetic distances between individuals showed two main groups, one including Kurils and Kamchatka (Pacific side group), and one including Newfoundland, Barents and White Seas (Atlantic side group, Figure 5). Atlantic side group showed more compact genetic distances between individuals than found in the Pacific side group. Atlantic side group also showed more star and chaplet topological subgraphs, commonly observed in clonal lineages (Stoeckel et al., 2024). In accordance with *ρST*, juveniles were more centrally connected in this network and less genetically distant to the other populations than the adults within each site. Some pairwise genetic distances between individuals within site, like in Kamchatka, were larger than between two individuals from different sites (Figure 5, e.g. Kamchatka’s individuals linked to White and Barents seas individuals).

**Figure 5:**
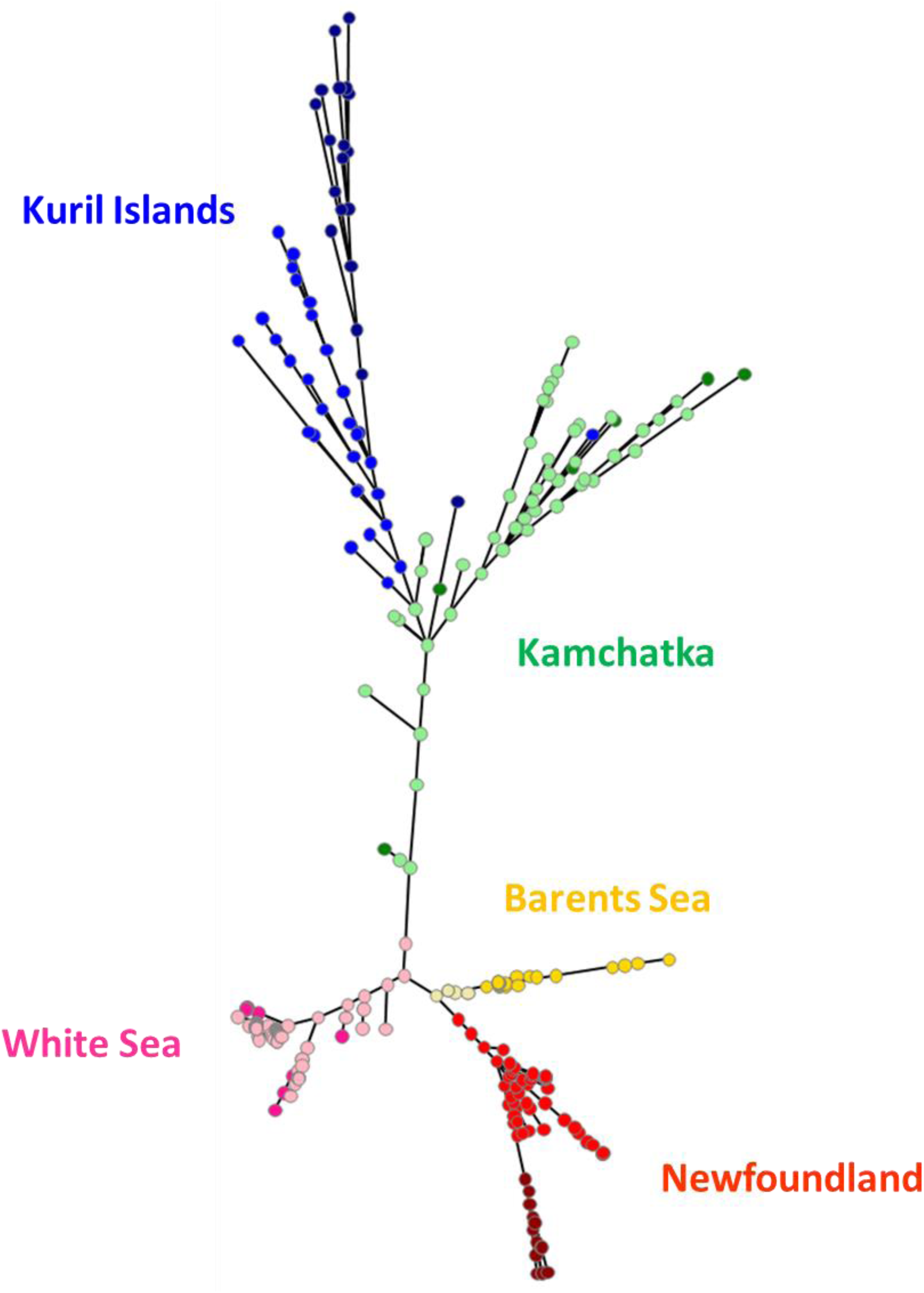
Minimum spanning tree of the genetic differentiation between sampled sites and cohorts of *A.stella* using pairwise *p*_*ST*_ using the ten nuclear microsatellites. Kamchatka adults and juveniles respectively in green and pale green; Kuril adults and juveniles respectively in dark blue and blue; Barents sea adults and juveniles respectively in gold and pale gold colors; White sea adults and juveniles respectively in pink and pale pink; Newfoundland adults and juveniles respectively in dark red and red.

## Discussion

We investigated reproductive modes, genetic diversity and differentiation in five populations of the autotetraploid sea anemone *Aulactinia stella*, sampled from North Pacific, Arctic, and North Atlantic coasts using three mitochondrial markers and ten nuclear microsatellites. *A. stella* is broadly distributed throughout the northern hemisphere where it is an important semi-sessile predator in shallow-water communities and prey for higher trophic levels (Ottaway, 1977). We genotyped two successive age cohorts from each of the five sites. We benefited from past sampling campaigns (from 2012 to 2017) to genotype individuals, with fewer sites and greater temporal dispersion in the Pacific Ocean: This could conceivably influence our results. The ten nuclear microsatellites we used offered sufficient power to distinguish clones from inbred offspring. Despite our standardized and reproducible protocol for allele dosage inference (Figure S2), tetraploid microsatellite datasets remain sensitive to technical biases, particularly those arising from null alleles and amplification biases, which may affect quantitative estimates prior to downstream analyses.

Genetic differentiation in *A. stella* revealed a hierarchical structure: first separating into a Pacific and an Arctic-Atlantic group, each of which was further subdivided by sampling site, and then by age cohort. This type of structure closely resembles patterns seen in the intertidal macroalga *Fucus distichus* and in other Arctic and peri-Arctic sessile coastal marine species (Coyer et al., 2011; Bringloe et al., 2020). Environmental factors, rather than species-specific biological or behavioral traits, may thus likely be the primary drivers of genetic differentiation in this region. We therefore examined reproductive modes and assessed genetic diversity and differentiation across these hierarchical groups.

### Reproductive modes in *Aulactinia stella* populations

We found strong genetic evidence that *A. stella* reproduces clonally in natural populations, consistent with the absence of males in some sites and with previous observations of embryos developing parthenogenetically within parental mesenteries (Bocharova, 2012, 2015; Kaliszewicz et al., 2012). In the North Pacific, repeated multilocus genotypes (MLGs) were rare; however, genetic indices, including interlocus FIS variances close to 1 and low levels of linkage disequilibrium, indicated intermediate levels of clonality (Navascues et al., 2010; Stoeckel et al., 2021a,b, 2024). It was further supported by changes in genotype frequencies between adults and juveniles, which yielded clonality estimates of 0.37 to 0.65 (Becheler et al., 2017; Stoeckel et al., 2024). The high clonal evenness in these populations likely reduced the probability of resampling identical MLGs (Arnaud-Haond et al., 2007, Stoeckel et al., 2021a). In contrast, Arctic and North Atlantic populations, characterized by lower allelic diversity, exhibited extensive clonality. Many individuals shared identical MLGs, and population-level genetic indices, including negative mean FIS, large variance of FIS across loci, and significant linkage disequilibrium, indicated high rates of clonal reproduction (Halkett et al., 2005; De Meeûs et al., 2006; Navascues et al., 2010; Stoeckel et al., 2021a). Consistently, clonality rates inferred from changes in genotype frequencies between adults and juveniles ranged from 0.74 to 1.0 in Arctic and North Atlantic populations, nearly twice the rates estimated for the North Pacific.

### The advantages of clonal reproduction at the limits of species’ ranges

High levels of uniparental reproduction, including clonality, are commonly observed in species expanding into new or marginal environments (Barrett, 2015; Pannel et al., 2015; Burke & Bonduriansky, 2019, Pereyra et al., 2023), particularly when enduring harsh conditions (Klimeš 2008) and when heading towards the poles (Peck et al., 1998). The post-glacial Baltic sea provides several examples, including the emblematic case of *Fucus vesiculosus* (Johannesson & André 2006; Pereyra et al., 2009; 2023). In plant species, the persistence of connection between ramets (mainly by roots) offers ecological advantages that may alone explain the prevalence of clonality at range margins, including regeneration after injury, distribution of physiological functions among ramets, translocation of resources between interconnected ramets (clonal integration) and space monopolization using different strategies (including phalanx versus guerrilla) to address intra- and inter-specific competitions depending on the environmental context (Cadotte et al., 2006; Gonzalvez et al., 2008; Roiloa, 2019; Franklin et al., 2021).

In contrast, dispersed clones in marine species such as *A. stella* do not benefit from such clonal integration. In this context, clonality should only favor kin cooperation (e.g., through brooding or cooperative behavior Bocharova, 2012; Rudin & Briffa, 2012) but more importantly it should favor the persistence of local populations under harsh conditions by avoiding the energetic and biological costs of sexual reproduction (Klimeš, 2008; Pierre et al., 2022).

Baker (1955) postulated that uniparental reproduction (including clonality) provides a critical advantage for colonization: individuals arriving in new areas can reproduce clonally to establish a new area. First, clonality allows a population to maintain itself locally until compatible mates become available (Baker & Stebbins, 1965, Daly et al., 2023). This hypothesis is particularly relevant for gonochoristic (or dioecious) sessile organisms that require close proximity for successful sexual reproduction (Barrett, 2015), such as in sea anemones. Second, in new harsh environments, clonal individuals that manage to survive minimize the risks of genetic reassortment associated with sexual reproduction for their offspring (Peniston et al., 2021; Pierre et al., 2022). Third, clonality maintains higher effective population sizes than in identical more sexual populations (Reichel et al. 2016; Stoeckel et al., 2021b). Clonality limits the negative effects of genetic drift during founder events, including the fixation of deleterious alleles in small populations (Navascues et al., 2010; He et al., 2024).

Our results in *A. stella*, which indicated a twofold increase in clonality at the species’ range edge, suggest that clonal reproduction may play an important role in supporting population persistence and maintaining higher effective population sizes and the more adapted genotypes under unfavorable conditions during range expansion (Baker’s conjecture, Pannel et al., 2015; Rafajlović et al., 2017; geographic parthenogenesis, Peck et al. 1998), without necessarily relying on mechanisms such as clonal integration (as proposed in plants) or habitat modification (as suggested for ecological engineering species) to explain clonal success.

### *Aulactinia stella* spreading across the Arctic Ocean from north Pacific?

Genetic diversity was nearly twice as high in North Pacific populations compared to Atlantic ones. Kamchatka and Kuril Islands harbored the greatest allelic and genotypic diversity, whereas Newfoundland, Barents Sea, and White Sea populations showed more compact identities in state. All Arctic and North Atlantic individuals carried the Pacific-3 haplotype, rare in the Pacific, while Kuril populations were fixed for Pacific-1, the dominant haplotype in Kamchatka. This distribution, considering our samples, suggests a North Pacific origin, with Kamchatka as a likely glacial refugium (He et al., 2024). In contrast, Arctic and North Atlantic populations may be more recently established (since the last glaciation) or alternatively, may have originated from smaller, more localized glacial refugia, evolving relatively isolated since the last warmer interglacial period (Eemian, minimum ∼116,000 years ago). But the DAPC results (Figure S4) and pairwise genetic distance (Figure 5) supported a stepping-stone genetic structure between sites and genetic groups rather than isolated as expected in an island model with no migration (Jombart et al., 2010). Interestingly, the path of the lowest *pST* genetic structure between sites align with major summer currents (e.g., Anadyr, Beaufort gyre, transpolar drift), areas of low ice cover, and coastal distances (Rudels & Carmack, 2022; Timmermans & Marshall, 2020; Constantin & Johnson, 2023) and increasing Arctic maritime traffic that may further accelerate dispersal, either through ballast water transport or passive enhancement by ship wakes (Melia et al., 2016; Boylan, 2021; PAME, 2024). Alterations in Arctic Ocean conditions, together with increasing human activities (including anthropogenic contributions to floating debris, ship-generated wake currents, and maritime transport of materials) may facilitate connectivity among *A. stella* populations, potentially via stepping-stone dispersal along coasts during the free-living polyp stage. Since at least 1993, temperatures in the upper 10 m of the water column have commonly reached 8–10 °C in coastal and sheltered bays, and 2–3 °C in the open sea, with higher temperatures in coastal and tidal currents along the Eurasian Arctic coasts, making this migration physiologically feasible (Locarnini et al. 2024; E.U. Copernicus Marine Service, 2024). These maritime and tidal currents are estimated to transport approximately 1 Sverdrup, a volume comparable to the combined mean discharge of all the world’s rivers, but substantially lower than the Gulf Stream, which rather transports 60–90 Sverdrups Constantin & Johnson, 2023). Taken together, these factors may explain the reduced genetic differentiation observed among juveniles within the established stepping-stone connectivity pattern of *A. stella* populations. Similar genetic patterns along with evidence for human-mediated dispersal have been reported for the coastal sea anemone *Diadumene lineata* (Glon et al., 2020), which, however, is more fecund and lacks the brooding behavior observed in *A. stella*.

Nevertheless, the biology of *A. stella*, like that of many Arctic marine organisms, remains poorly understood once individuals move beyond nearshore habitats and become inaccessible to direct observation. In particular, dedicated studies are needed to determine whether this Arctic sea anemone can persist while attached to debris or as free-living polyp or adult drifters within coastal and tidal currents, potentially enabling effective migration across the Arctic Ocean.

### Juveniles less genetically structured than adults: methodological artifacts or gene flow with local endemism?

Comparisons of genetic diversity between adults and juveniles revealed slightly lower levels of genetic differentiation among juveniles (Figure S3). Several non-exclusive hypotheses could explain this pattern, requiring further research to disentangle their relative contributions. Methodological artefacts must first be ruled out through more geographically balanced and systematically designed sampling. Using polymorphic markers with lower mutation rates could also help distinguish genuine shared ancestry from homoplasy that has higher chance to occur in microsatellite loci across distant populations.

Clarifying the life cycle of *A. stella*, particularly the dynamics of clonal interference and mortality during recruitment, will be essential to assess whether reduced genetic structure in juveniles and the absence of shared multilocus genotypes (MLGs) across sites reflect local genetic endemism, gregarious behavior promoting shared ancestry (as found in other sea anemone species: Lane et al., 2020; Ryan et al., 2021), or persistent founder effects with limited gene flow. *A. stella* occupies a range of habitats, such as the Barents and White Seas, which differ significantly in salinity despite their proximity (∼100 km). Environmental factors, including temperature, salinity, depth, tidal exposure, and substrate disturbance, are known to influence reproductive modes and genetic differentiation (Eriksson & Johansson, 2005; Krueger-Hadfield et al., 2013; Berković et al., 2014; Dudgeon et al., 2017; McMahon et al., 2017; Ryan & Miller, 2019; Williams et al., 2024). Similar habitat-driven patterns are observed in other sea anemones, such as *Actinia tenebrosa* and *Diadumene lineata*, where localized clonal distributions and endemic lineages correlate with temperature, salinity, UV exposure, and community composition (Sherman & Ayre, 2008; Ryan et al., 2021). It may explain the higher genetic differentiation found in adults between sites.

### Restricted dispersal or local adaptation in *Aulactinia stella*?

No MLGs were shared between sampling sites, even over relatively shorter distances (e.g., Barents vs. White Sea, ∼360 km; Kuril Islands vs. Kamchatka, ∼680 km). The presence of local, site-specific clones and high levels of genetic differentiation between adult populations is common in cold-water sea anemones, including *A. tenebrosa*, *Anthothoe albocincta*, *Diadumene lineata*, and several North Pacific species (Veale & Lavery, 2012; Sherman et al., 2007; Ryan et al., 2021; Edmands & Potts, 1997), which contrasts with tropical species that often share genets across hundreds of kilometers (Russo et al., 1994; Emms et al., 2020). Such patterns would suggest that Arctic conditions may constrain and challenge the dispersal of sea anemones (Chan et al., 2019), but other passively dispersing taxa in cold-water habitats such as macroalgae, seagrasses, and bivalves, show weaker population structure (Coolen et al., 2020; Pereyra et al., 2023; Krueger-Hadfield et al., 2021). In fact, polyp stage commonly allows locally-active and passive transport by tidal flows and circumpolar maritime currents, facilitating long-distance dispersal (Riemann-Zürneck, 1998). For instance, *Urticina eques* exhibits very low differentiation over hundreds of kilometers along the British coast (Cava-Solé et al., 1994). In the case of *A. stella*, Arctic populations may have only narrow temporal windows of suitable thermal conditions for successful sexual reproduction, particularly in the intertidal and shallow sublittoral zones. These habitats are also characterized by extreme variability in salinity and desiccation at low tide. In such habitats, *A. stella* is often found hiding under *Fucus* algae, in rock crevices, or in tide pools, behaviors that indicate exposure to environmental stress. Local environmental filtering through selection and post-settlement mortality (e.g., *Fucus vesiculosus*: Johannesson & André, 2006; Pereyra et al., 2009; *Fucus distichus*: Laughinghouse et al., 2015; *Actinia tenebrosa*: Sherman et al., 2008; seagrasses: Kendrick et al., 2012; *Acropora millepora*: van Oppen et al., 2011), as well as individual habitat preferences and gregarious settlement (e.g., *Diadumene lineata*: Ryan et al., 2021), may explain the roughly twofold higher genetic differentiation observed among adults compared to juveniles in *A. stella*. These patterns highlight the need for further ecological and behavioral studies to clarify the potential mechanisms and temporal dynamics of dispersal in this species in the Arctic Ocean.

## Data Availability

All data (including adults, juveniles, MLG genotypes, sampling GPS locations, and clone sizes) as well as the scripts used to reproduce the results and figures, are openly available on Zenodo https://doi.org/10.5281/zenodo.17193174

## Supplementary Material

Supplementary material provides two tables and three figures. Table S1 reports distributions of repeated MLGs within sites and MLGs shared between adults and juveniles; Table S2 reports pairwise *pST* between sites and between age cohorts; Figure S1 reports a map of the sample sites with the minimum spanning trees of genetic differentiation of adults, juveniles and overall among sites; Figure S2 reports how allele dosage of the microsatellites was achieved; Figure S3 reports the change of the Bayesian information criteria of the DAPC with increasing number of genetic clusters arguing for a stepping-stone genetic structure.

## Acknowledgements

The authors thank N.P. Sanamyan, K.E. Sanamyan, B.V. Osadchenko and I.A. Kosevich for providing material and assistance in species identifications, N.S. Mugue, T.V. Neretina, A.E. Barmintseva and S.D. Grebelnyi for professional consultations, A. Mercier, J.-F. Hamel, E. Montgomery and the Ocean Sciences Field Services of Memorial University for their assistance with the animal collections. We thank Patricia Nadan and Pascale Le Névé (INRAE IGEPP) for their administrative support along this project.

A preprint version of this article has been peer-reviewed and recommended by PCI Evol Biol (https://doi.org/10.24072/pci.evolbiol.100866). We warmly thank Benoit Nabholz, Kerstin Johannesson, Malgorzata Lagisz and one anonymous reviewer for their insightful comments that helped improving our manuscript and its data availability and reproducibility.

## Author contributions

EB and SS laid the foundation of this work while EB visited SS lab in 2018 and 2019. EB and SS conceived the study and were responsible for funding applications. EB was responsible for data acquisition. She supervised and achieved sample collection, determined the species and sexed the sampled individuals. She also conceived, developed the sequences and microsatellites, supervised and achieved sequencing and genotyping of all the samples. From sequencing machines, she genotyped the samples. AV contributed to methodology, supervision and administration of data acquisition. EB performed data analyses and produced figure on mitochondrial sequences. SS performed data analyses and produced tables and figures on microsatellites. SS and EB tracked and managed the bibliography, interpreted results, wrote the manuscript and managed its edition. All authors read and approved this manuscript.

## Supplementary information

**Figure S1:**
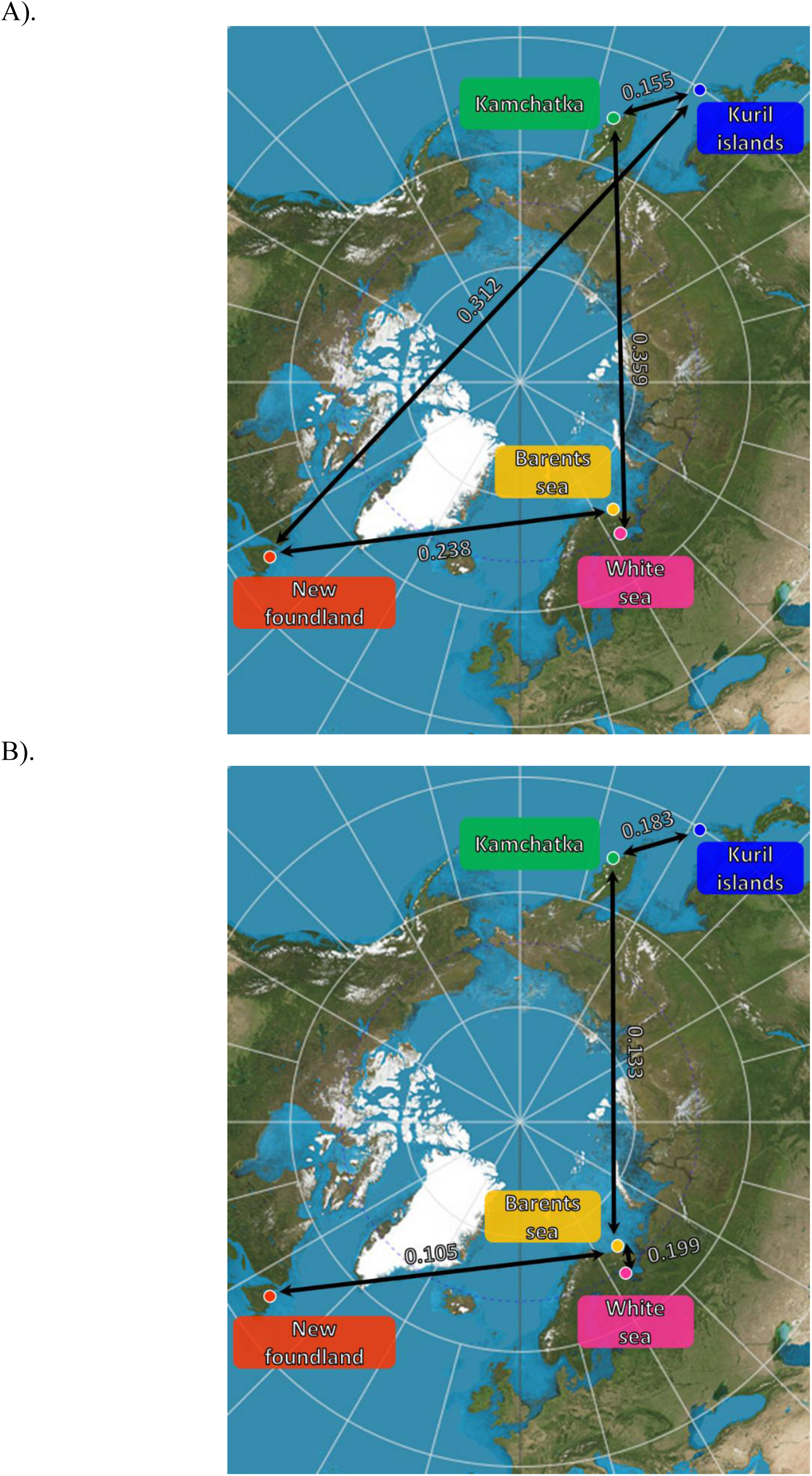
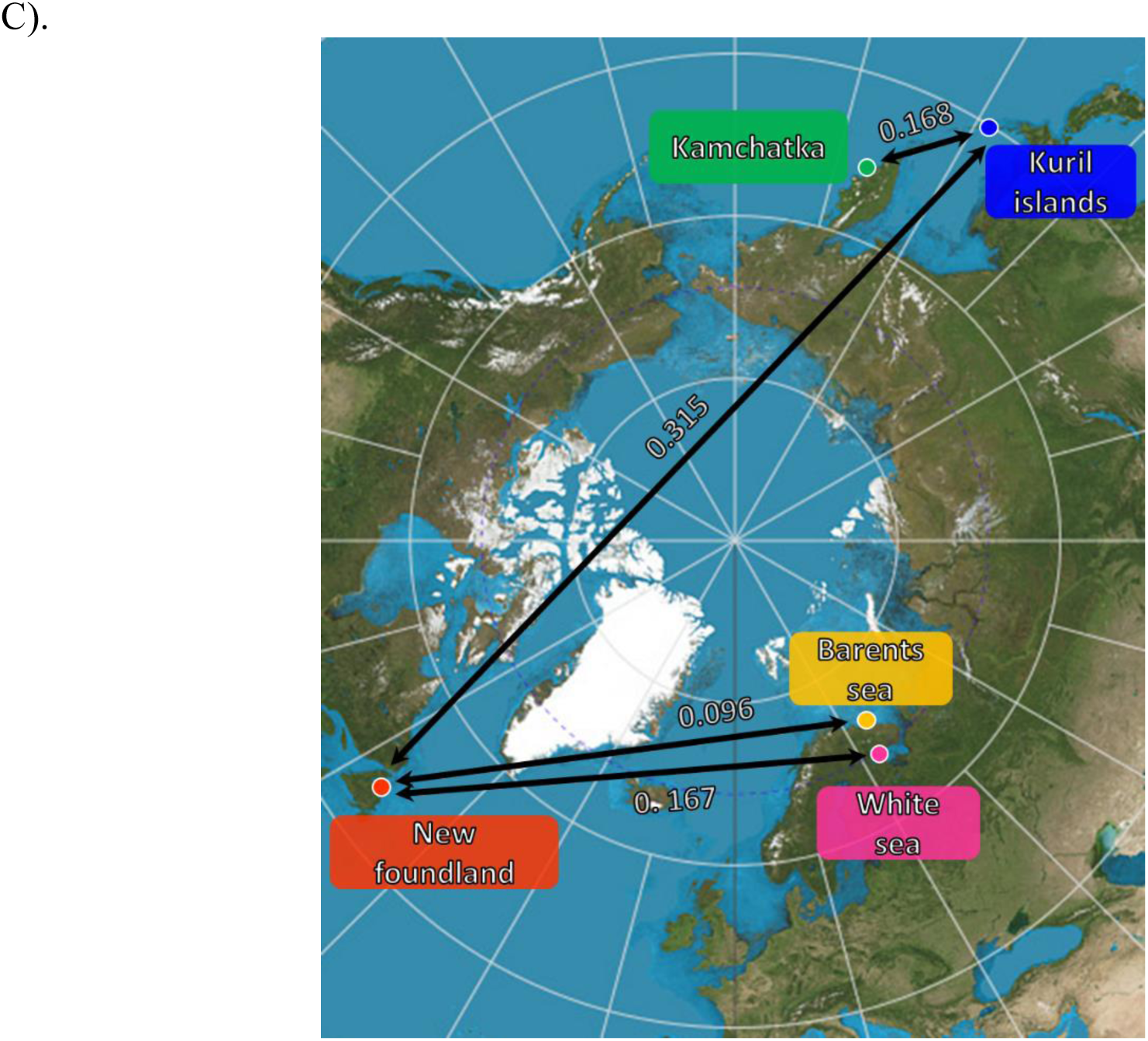
Minimum spanning tree of A). the *pST* of adults, B). the *pST* of juveniles and C). the *pST* of all age cohorts over all loci between sampled sites.

**Figure S2:**
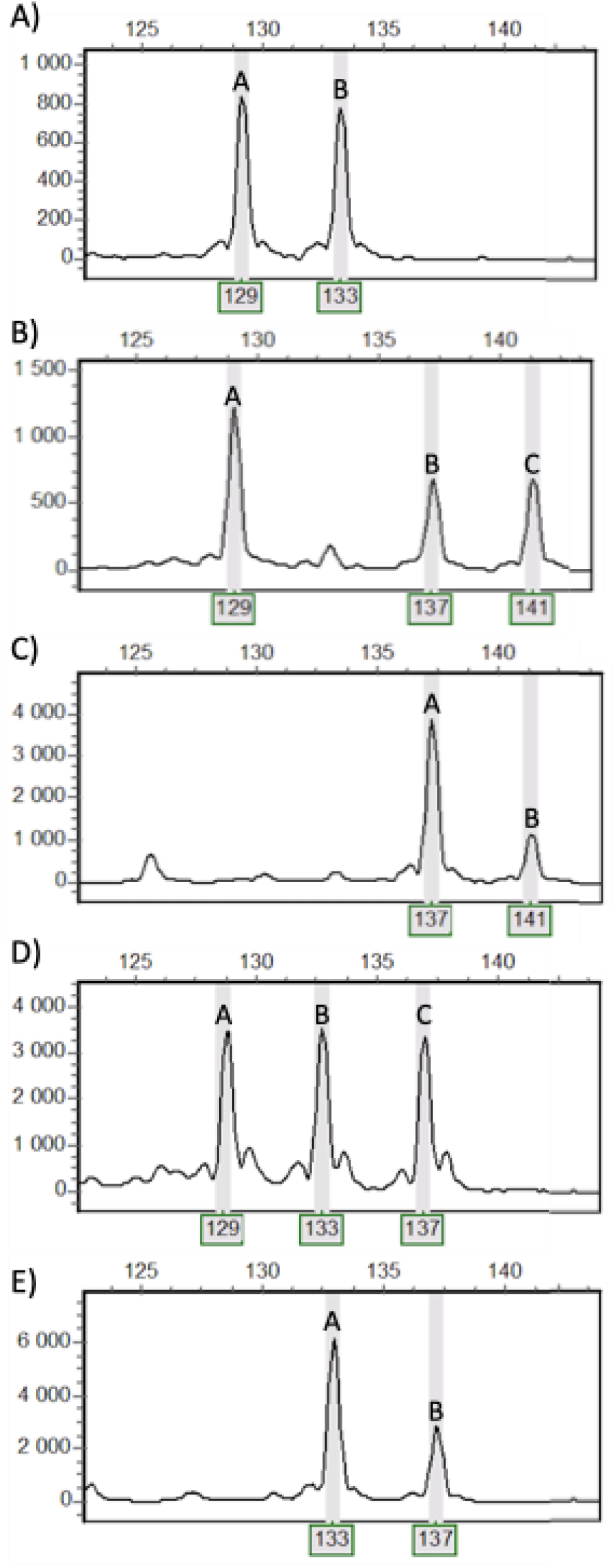
Counting of allele dosage of the microsatellite locus 249TAM (as an example) in *Aulactinia stella* (within each individual per locus): A). 129, 129, 133, 133 (AABB), B). 129, 129, 137, 141 (AABC), C). 137, 137, 137, 141 (AAAB), D). 129, 133, 137 (ABC+null allele), E). 133, 133, 137 (AAB+null allele).

**Figure S3:**
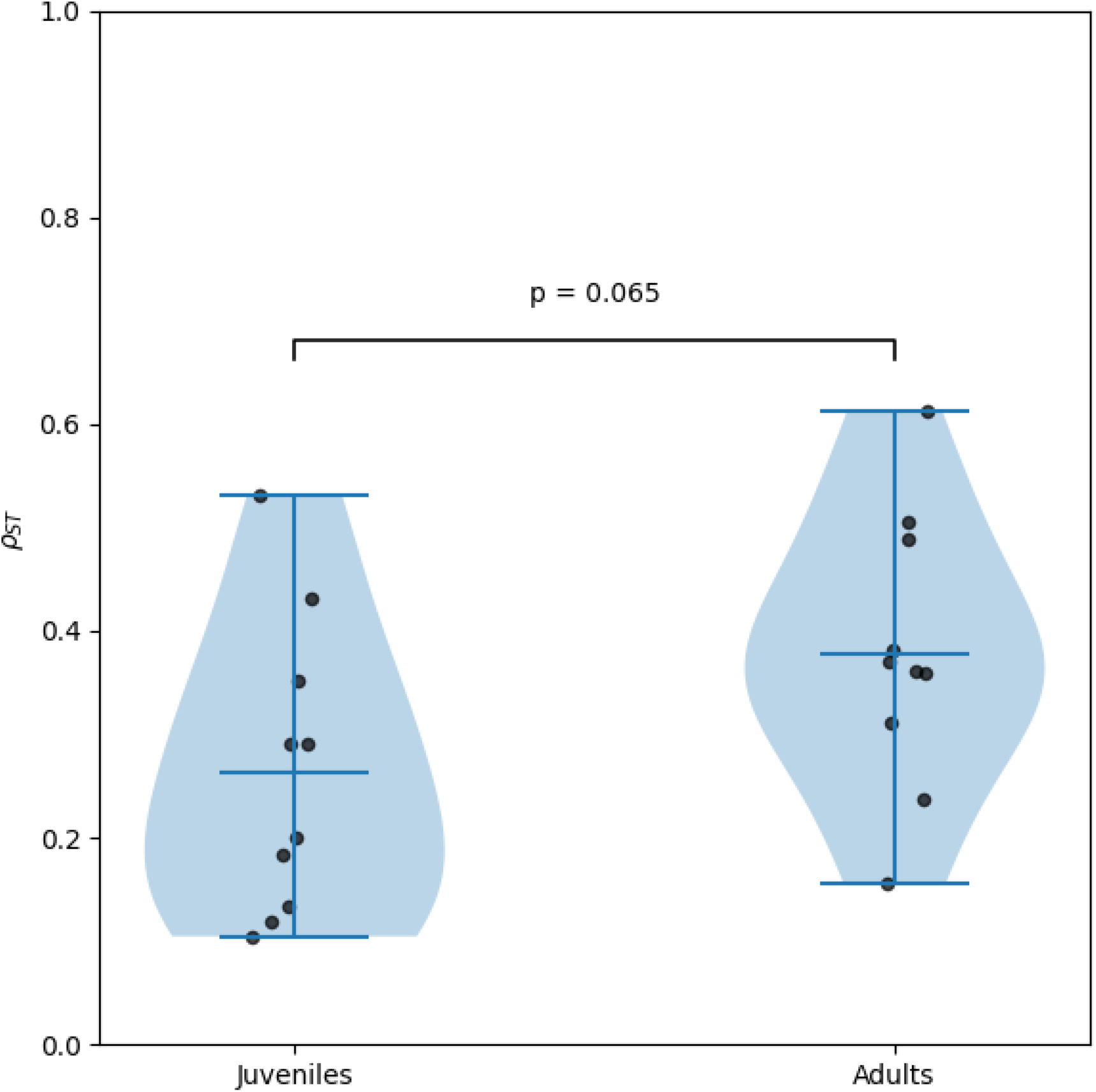
Distributions of *pST* between pairs of sites among juveniles and adults (light blue violin plots). We reported the probability that difference of *pST* between juveniles and adults would be due to a random (sampling, locus) effects as a Wilcoxon test (significance bar), the *pST* values in each age cohort (Black dots), the extremum and mean values (dark blue bars).

**Figure S4:**
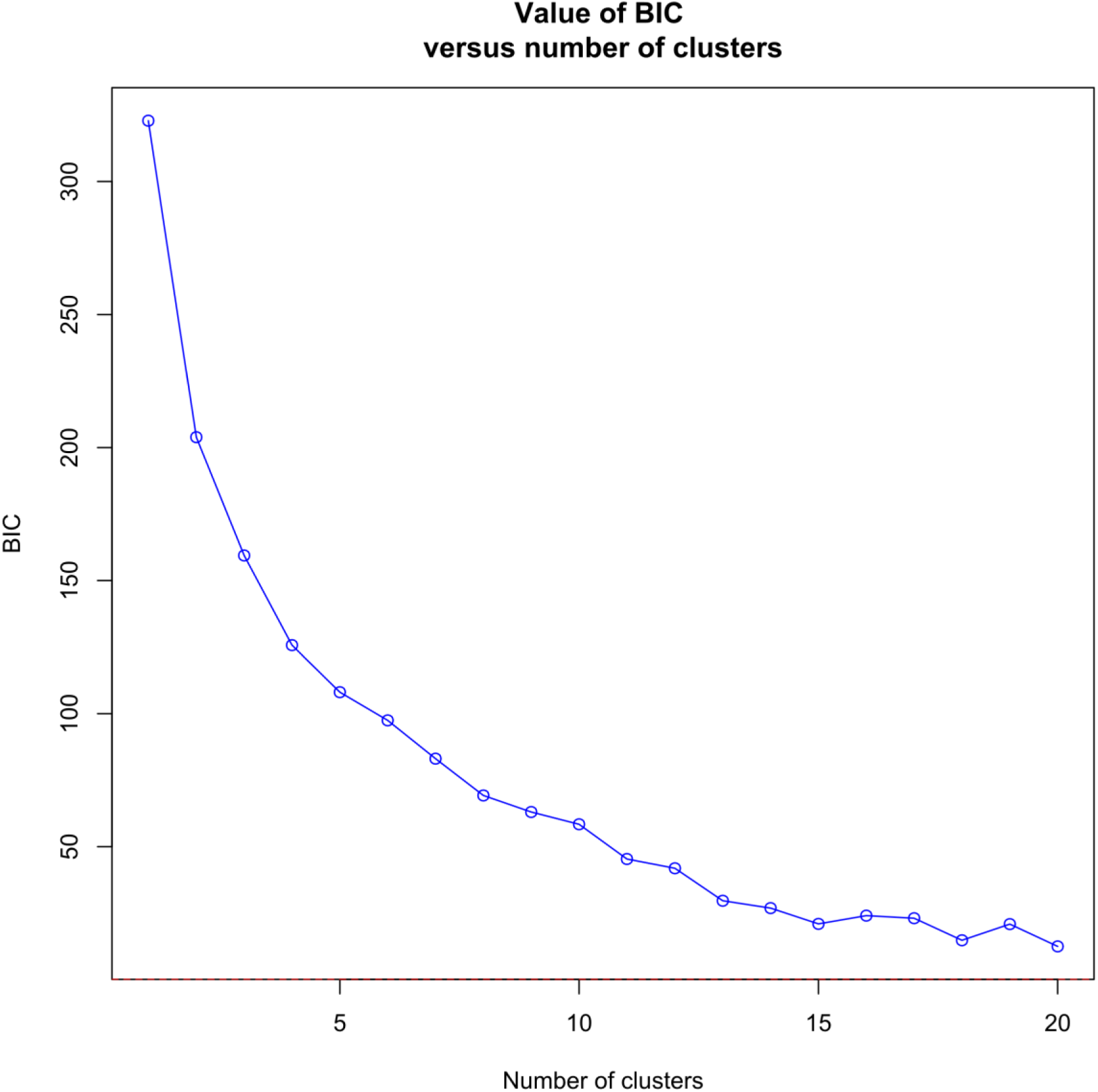
Change of the Bayesian information criteria (BIC) with increasing number of genetic clusters used in the Discriminant Analysis of Principal Components to investigate the genetic differentiation between individuals and among sampled sites and cohorts. The continuously decreasing pattern observed is typical of a stepping-stone structure of the studied demes.

**Table S1:**
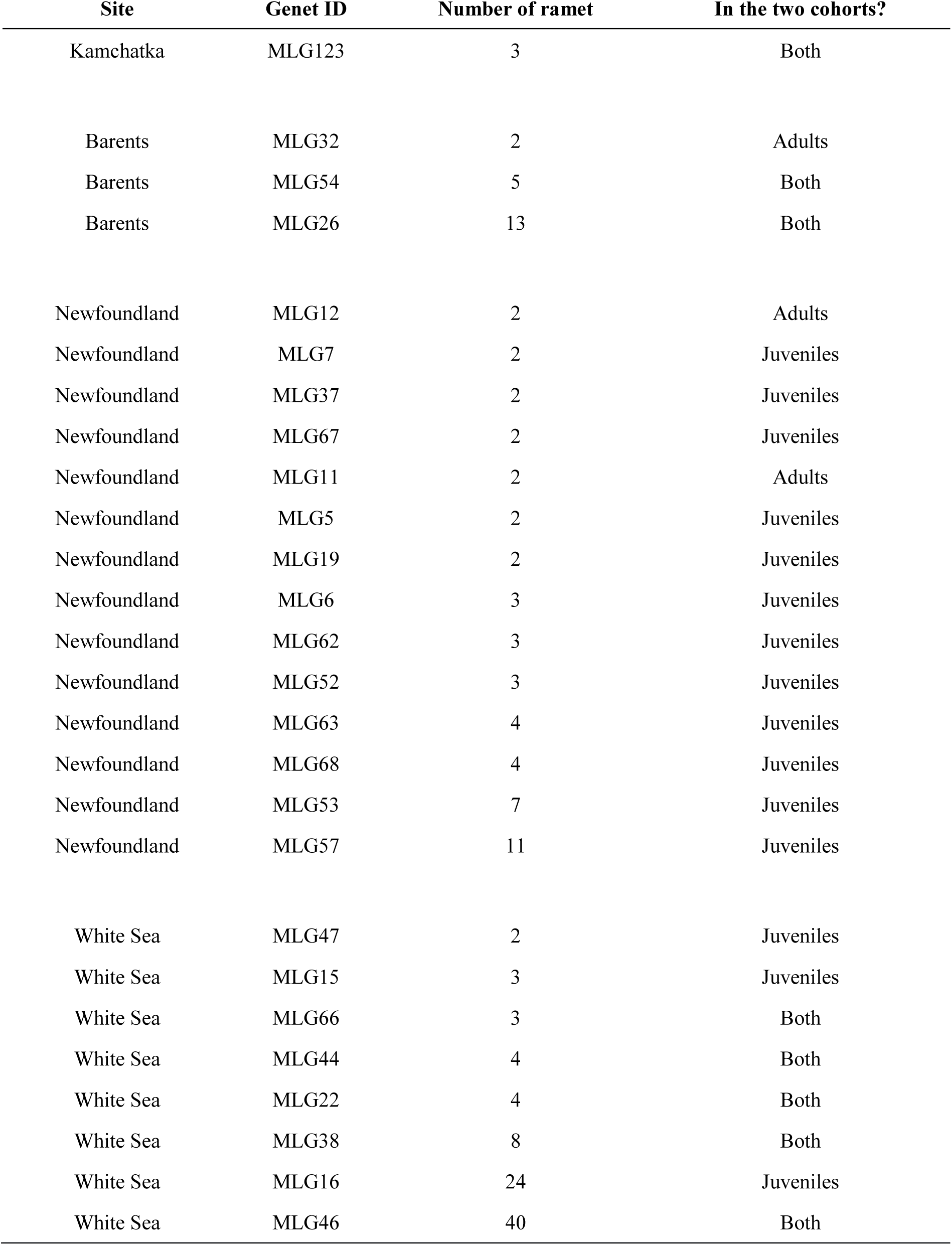
Distribution of repeated MLGs within sites and MLGs shared between adults and juveniles. Genet ID, the number of ramets within the genet and if the considered genet has been found in adults, juveniles or both are reported. No repeated MLG was found in Kuril islands sampled genotypes.

| Site | Genet ID | Number of ramet | In the two cohorts? |
| --- | --- | --- | --- |
| Kamchatka | MLG123 | 3 | Both |
| Barents | MLG32 | 2 | Adults |
| Barents | MLG54 | 5 | Both |
| Barents | MLG26 | 13 | Both |
| Newfoundland | MLG12 | 2 | Adults |
| Newfoundland | MLG7 | 2 | Juveniles |
| Newfoundland | MLG37 | 2 | Juveniles |
| Newfoundland | MLG67 | 2 | Juveniles |
| Newfoundland | MLG11 | 2 | Adults |
| Newfoundland | MLG5 | 2 | Juveniles |
| Newfoundland | MLG19 | 2 | Juveniles |
| Newfoundland | MLG6 | 3 | Juveniles |
| Newfoundland | MLG62 | 3 | Juveniles |
| Newfoundland | MLG52 | 3 | Juveniles |
| Newfoundland | MLG63 | 4 | Juveniles |
| Newfoundland | MLG68 | 4 | Juveniles |
| Newfoundland | MLG53 | 7 | Juveniles |
| Newfoundland | MLG57 | 11 | Juveniles |
| White Sea | MLG47 | 2 | Juveniles |
| White Sea | MLG15 | 3 | Juveniles |
| White Sea | MLG66 | 3 | Both |
| White Sea | MLG44 | 4 | Both |
| White Sea | MLG22 | 4 | Both |
| White Sea | MLG38 | 8 | Both |
| White Sea | MLG16 | 24 | Juveniles |
| White Sea | MLG46 | 40 | Both |

**Table S2:**
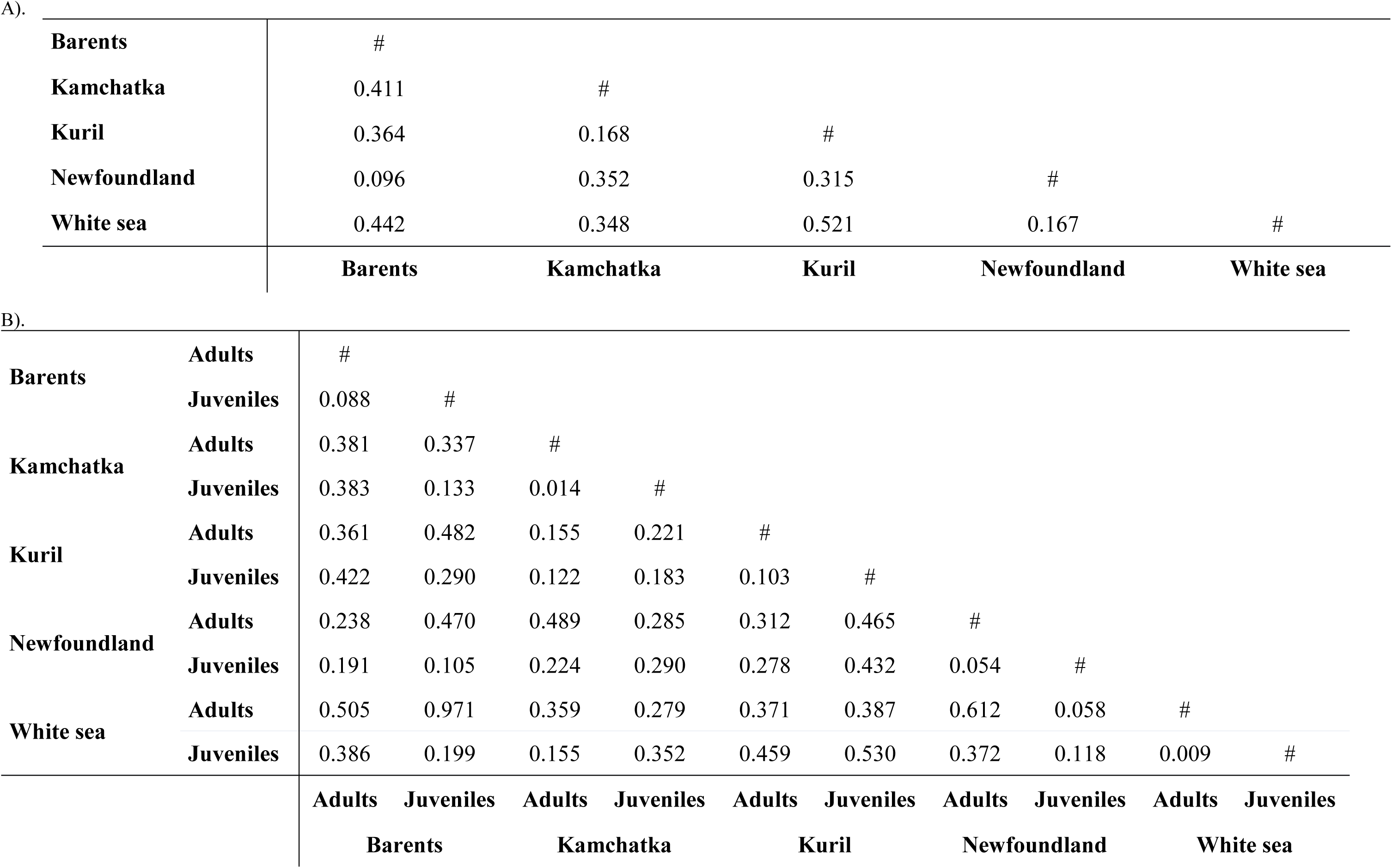
Pairwise *pST* between sites (A) and between age cohorts within sites and among sites (B).

|  |  |  |  |  |  |
| --- | --- | --- | --- | --- | --- |
| **Barents** | # |  |  |  |  |
| **Kamchatka** | 0.411 | # |  |  |  |
| **Kuril** | 0.364 | 0.168 | # |  |  |
| **Newfoundland** | 0.096 | 0.352 | 0.315 | # |  |
| **White sea** | 0.442 | 0.348 | 0.521 | 0.167 | # |
|  | **Barents** | **Kamchatka** | **Kuril** | **Newfoundland** | **White sea** |

**Table S2:** Pairwise $\rho_{ST}$ between sites (A) and between age cohorts within sites and among sites (B).
|  |  |  |  |  |  |  |  |  |  |  |  |
| --- | --- | --- | --- | --- | --- | --- | --- | --- | --- | --- | --- |
| **Barents** | **Adults** | # |  |  |  |  |  |  |  |  |  |
|  | **Juveniles** | 0.088 | # |  |  |  |  |  |  |  |  |
| **Kamchatka** | **Adults** | 0.381 | 0.337 | # |  |  |  |  |  |  |  |
|  | **Juveniles** | 0.383 | 0.133 | 0.014 | # |  |  |  |  |  |  |
| **Kuril** | **Adults** | 0.361 | 0.482 | 0.155 | 0.221 | # |  |  |  |  |  |
|  | **Juveniles** | 0.422 | 0.290 | 0.122 | 0.183 | 0.103 | # |  |  |  |  |
| **Newfoundland** | **Adults** | 0.238 | 0.470 | 0.489 | 0.285 | 0.312 | 0.465 | # |  |  |  |
|  | **Juveniles** | 0.191 | 0.105 | 0.224 | 0.290 | 0.278 | 0.432 | 0.054 | # |  |  |
| **White sea** | **Adults** | 0.505 | 0.971 | 0.359 | 0.279 | 0.371 | 0.387 | 0.612 | 0.058 | # |  |
|  | **Juveniles** | 0.386 | 0.199 | 0.155 | 0.352 | 0.459 | 0.530 | 0.372 | 0.118 | 0.009 | # |
|  |  | **Adults** | **Juveniles** | **Adults** | **Juveniles** | **Adults** | **Juveniles** | **Adults** | **Juveniles** | **Adults** | **Juveniles** |
|  |  | **Barents** |  | **Kamchatka** |  | **Kuril** |  | **Newfoundland** |  | **White sea** |  |

